# MMinte: An application for predicting metabolic interactions among the microbial species in a community

**DOI:** 10.1101/059550

**Authors:** Helena Mendes-Soares, Michael Mundy, Luis Mendes Soares, Nicholas Chia

**Author notes:** **<u>Helena Mendes-Soares</u>** 200 First St. SW Mayo Clinic Rochester, MN 55905 USA <u>Michael Mundy</u> Mayo Clinic USA **<u>Luis Mendes Soares</u>** Harvard Medical School USA **<u>Nicholas Chia</u>** Mayo Clinic USA.

## Abstract

**Background.:** The explosive growth of microbiome research has yielded great quantities of data. These data provide us with many answers, but raise just as many questions. 16S rDNA—the backbone of microbiome analyses—allows us to assess α-diversity, β-diversity, and microbe-microbe associations, which characterize the overall properties of an ecosystem. However, we are still unable to use 16S rDNA data to directly assess the microbe-microbe and microbe-environment interactions that determine that system's broader ecology. Thus, properties such as competition, cooperation, and nutrient conditions remain insufficiently analyzed. Here, we apply predictive community metabolic models of microbes identified with 16S rDNA data to probe the ecology of microbial communities.

**Results.:** We developed a methodology for the large-scale assessment of microbial metabolic interactions (MMinte)from 16S rDNA data. MMinte assesses the relative growth rates of interacting pairs of organisms within a community metabolic network and whether that interaction has a positive or negative effect. Moreover, MMinte's simulations take into account the nutritional environment, which play a strong role in determining the metabolism of individual microbes. We present two case studies that demonstrate this software's utility. In the first, we show how diet influences the nature of the microbe-microbe interactions. In the second, we use MMinte's modular feature set to better understand how the growth of *Desulfovibrio piger* is affected by, and affects the growth of, other members in a simplified gut community under metabolic conditions suggested to be determinant for their dynamics.

**Conclusion.:** By applying metabolic models to commonly available sequence data, MMinte grants the user insight into the metabolic relationships between microbes, highlighting important features that may relate to ecological stability, susceptibility, and cross-feeding. These relationships are at the foundation of a wide range of ecological questions that impact our ability to understand problems such as microbially-derived toxicity in colon cancer.

## Background

Advances in sequencing technology have culminated in an explosion of 16S rDNA-based microbiome projects, both small [1] [2] [3] and large [4, 5] [6], The microbial ecosystems characterized in these projects are the basis for many critical life processes, from global nutrient cycles [7, 8] to homeostasis in the human body [9–11]. The importance of microbiome research is embodied in recent calls for the formation of a worldwide microbiome consortium [12]. The gut microbiome exemplifies a complex system and contains trillions of interacting bacterial cells [13]. It is not sufficient to treat bacterial taxa as independent entities in a statistical framework of association and diversity. Instead, ecological investigation requires examining the biological interactions underlying the complexities of our microbial communities [14].

Efforts to understand complex microbial communities range from inference based on 16S rDNA sequences [15] to the use of ‘omics technologies across multiple time points [16]. A variety of software and tools for analyzing 16S rDNA data exist, and range from identifying taxa [17, 18] and calculating diversity [19] to producing microbe-microbe association networks [20] However, none of these utilize 16S rDNA to understand the mechanistic basis of microbe-microbe interactions. Each measure captures part of a complex picture, but none captures the functional basis [21] for the microbial interactions that make up a community—i.e., the building blocks of the microbiome.

Bridging the gap between association and mechanism in microbe-microbe interactions requires an approach centered on mechanistic principles. One avenue to deciphering the role of a microbe in a community is through the use of a predictive modeling approach [22, 23]. Metabolic models recapitulate the biological processes of nutrient uptake and metabolite secretion [24], which are at the basis of most microbial interactions. Computationally, the reconstruction of genome-scale metabolic models [25, 26] has been automated through large-scale computing efforts such as RAST [27] and ModelSEED [28]. Tools such as COBRA Toolbox [29, 30] provide an interface for manipulating and investigating metabolic network models. Recently, community metabolic models have been generated to explore the gut microbiome in health and disease [31–34], but these efforts have been driven largely by manual curation—a time consuming and laborious practice [26]. Building on these past research efforts, we explore an alternative path to generating predictive community metabolic models for large-scale microbial communities.

MMinte (pronounced /'minti/) is an integrated pipeline that allows users to explore the pairwise interactions (positive or negative) that occur in a microbial network. From an association network and 16S rDNA sequence data, MMinte identifies corresponding genomes, reconstructs metabolic models, estimates growth under specific metabolic conditions, analyzes pairwise interactions, assigns interaction types [35] to network links, and generates the corresponding network of interactions. Our application is composed of a set of seven individual functionalities, known as widgets, that run sequentially, and each widget may also be run as an independent module. Below, we present two case studies from the gut microbiome that illustrate how MMinte can be used to predict ecological features of a microbial community based on metabolic maps of bacterial species. In doing so, MMinte provides a valuable tool for generating well-defined mechanistic hypotheses for further exploration.

## Implementation

MMinte consists of minimally overlapping functions that come together to perform a single task. In designing it, our goal was to facilitate code re-use by focusing on modularity, allowing the user to streamline the parts presented here for other purposes. Indeed, we do not view MMinte as a single-purpose code, but as a set of widgets that can be repurposed for multiple queries, ranging from testing interactions between a set pair of species to reconstructing a community metabolic network.

The web browser interface creates a point-and-click experience that allows the user to perform complex analysis on large data sets without programming expertise. Forthose seeking more control orto implement their own pipelines using MMinte widgets, MMinte functions can also be run in a command-line environment. Because all of the code is provided to the user, it can be changed to fit a particular need. Finally, MMinte is under continuous development, it is publicly available on Github (http.github.com/mendessoares/MMinte) for use by the community, and the authors welcome contributions to further its development.

A full run of MMinte generates a predicted network of microbe-microbe interactions for a microbial community using a sequence of seven widgets that progressively analyze 16S rDNA sequences, then genomes, metabolic models, and finally community metabolic networks. The analysis can be run uninterrupted, and all intermediate files are stored. The seven widgets that constitute MMinte are depicted in **Figure 1**.

**Figure 1.**
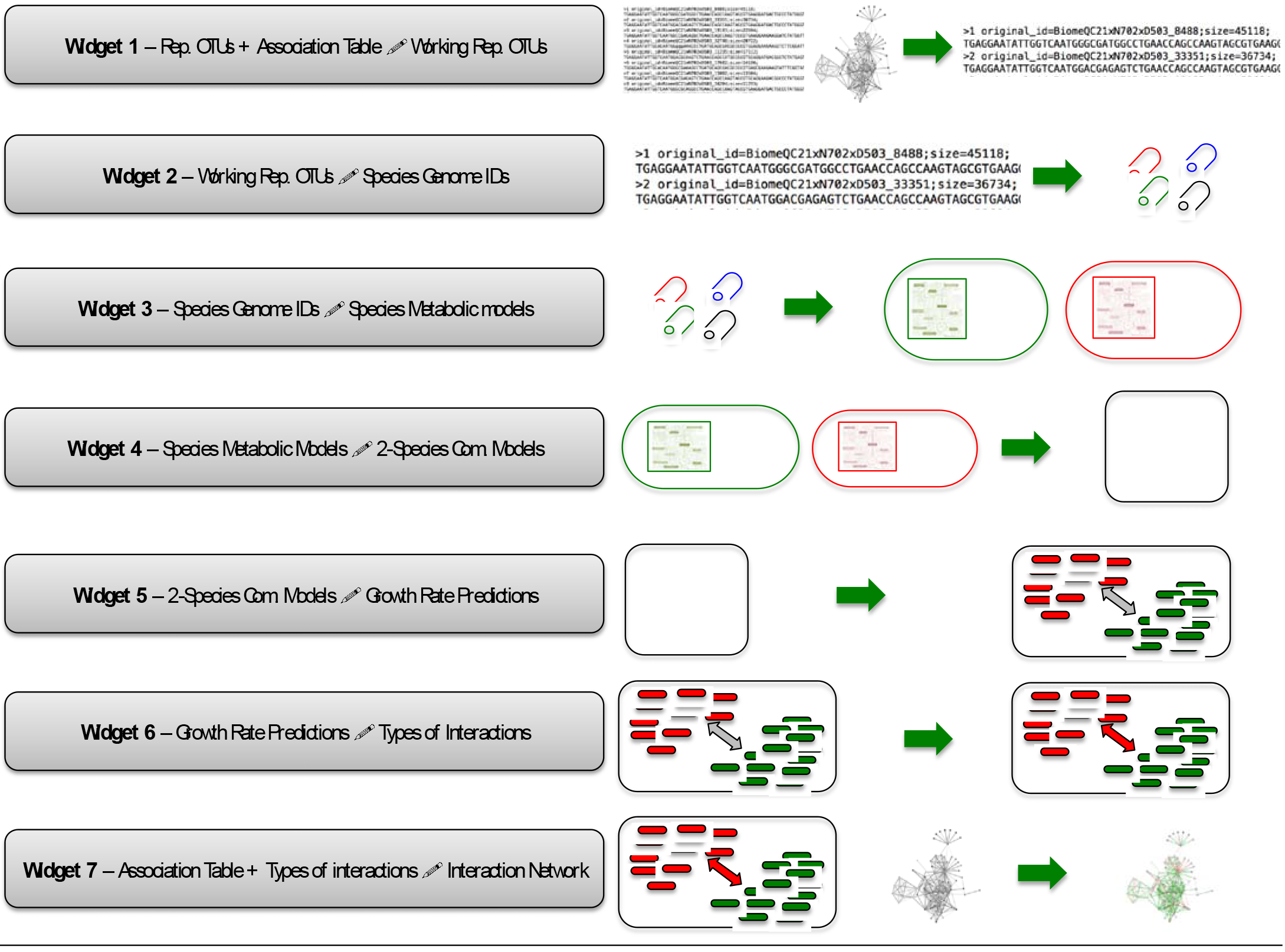
Schematic of the MMinte pipeline. Each green rectangle represents one widget. **Widget 1** takes two files, a network of associations between operational taxonomic units (OTUs) and a FASTA file containing the 16S rDNA sequences from a microbiome study, and reduces the latter data set to include only the sequences for OTUs present in the network. **Widget 2** identifies the sequences provided and assigns them a genome ID. The percent similarity between the query OTU and the 16S sequence of the genome to which it was matched is stored in a file to be used by **Widget 7. Widget 3** Calls the ModelSEED service [28] with the list of genome IDs produced by Widget 2, which reconstructs species metabolic models that are exported to the user's local machine. **Widget 4** then uses these species models to create metabolic models for two-species communities. **Widget 5** estimates the growth rate of each species in the community under defined metabolic conditions, which can be changed by the user. **Widget 6** assesses the types of interaction (mutualism, parasitism, commensalism, competition, amensalism, or neutralism) occurring between the pairs of species in a community based on the effect that each species has on the growth of another. **Widget 7** takes the initial information about the topology of the network, the information about the percent similarity between OTUs and the closest genomes, and the types of interactions and plots an interaction network in which the color of the links represents the type of interaction (positive, green; negative, red; no interaction, grey).

<u>Widget 1:</u> *Reduces data for the downstream analysis*. The purpose of this step is to remove operational taxonomic units (OTUs) that will not be used in future analyses, based on an existing list of OTU associations.

*Inputs*: (1) 16S rDNA sequences of all representative OTUs (as one might obtain from, for instance, QIIME [36] or mothur [37]) and (2) an association table between pairs of OTUs.

*Output*: 16S rDNA sequences of OTUs.

<u>Widget 2</u>: *Matches 16S rDNA signatures with corresponding genomes*. Using BLAST, the user's 16S rDNA sequences are matched with 16S rDNA sequences from publicly available, complete genomes in NCBI [38]. Importantly, we output a table of percent similarities between OTU-and genome-derived 16S rDNA sequences. This information is used to limit potential sources of error from imperfect OTU-genome pairings and to color code nodes in the final network (*Widget7*).

*Input*: 16S rDNA sequences of OTUs.

*Output*: (1) Genome IDs and (2) percent similarity table.

<u>Widget 3</u>: *Obtains metabolic models*. This widget takes advantage of the ModelSEED [28] framework to reconstruct and gap-fill metabolic models for a list of genomes.

*Input*: Genome IDs.

*Output*: Single-species metabolic models.

<u>Widget 4</u> *Merges models*. Using COBRApy [39], this function creates metabolic models of two-species communities from a list of pairs of species [33, 40]. The list can be provided by the user or created by MMinte.

*Input*: (1) Species-species associations (optional) and (2) single-species metabolic models in the Systems Biology Markup Language (SBML) format.

*Output*: Two-species metabolic models.

<u>Widget 5</u> *Runs flux balance analysis*. This step estimates the growth rates for each species under defined nutrient conditions, in isolation and in the presence of another species, by running a flux balance analysis in COBRApy [39, 41]. The user has the option to specify the nutrient conditions to reflect the specific conditions of the environment being studied.

*Inputs*: (1) Two-species metabolic models and (2) choice of metabolic conditions to be used from media file (provided in the supportFiles folder, default choice = “complete”).

*Output*: Growth-rate predictions.

<u>Widget 6</u> *Evaluates metabolic interactions*. Using previously calculated growth rates, this function quantifies the effect of pairwise interactions and assigns an interaction type to each pair, following Heinken and Thiele [33]. The possible types are positive (commensalism, mutualism), negative (parasitism, amensalism, competition), or no (neutralism) interaction.

*Input*: Growth-rate predictions.

*Output*: Quantitative effect of interaction and interaction type predicted.

<u>Widget 7</u> *Draws community metabolic network*. This function generates a color-coded interaction network using the D3.js [42] visualization platform, based on the associations provided to Widget 1. Links are colored according to the type of interaction predicted by MMinte (*Widget 6*). A node's shading reflects the percent similarity between OTUs and genomes (*Widget 2*)

*Input*: (1) Association table between pairs of OTUs, (2) percent similarity table, and (3) quantitative effect of interaction and interaction type predicted.

*Output*: Metabolic interaction network (see **Fig. 2**)

**Figure 2.**
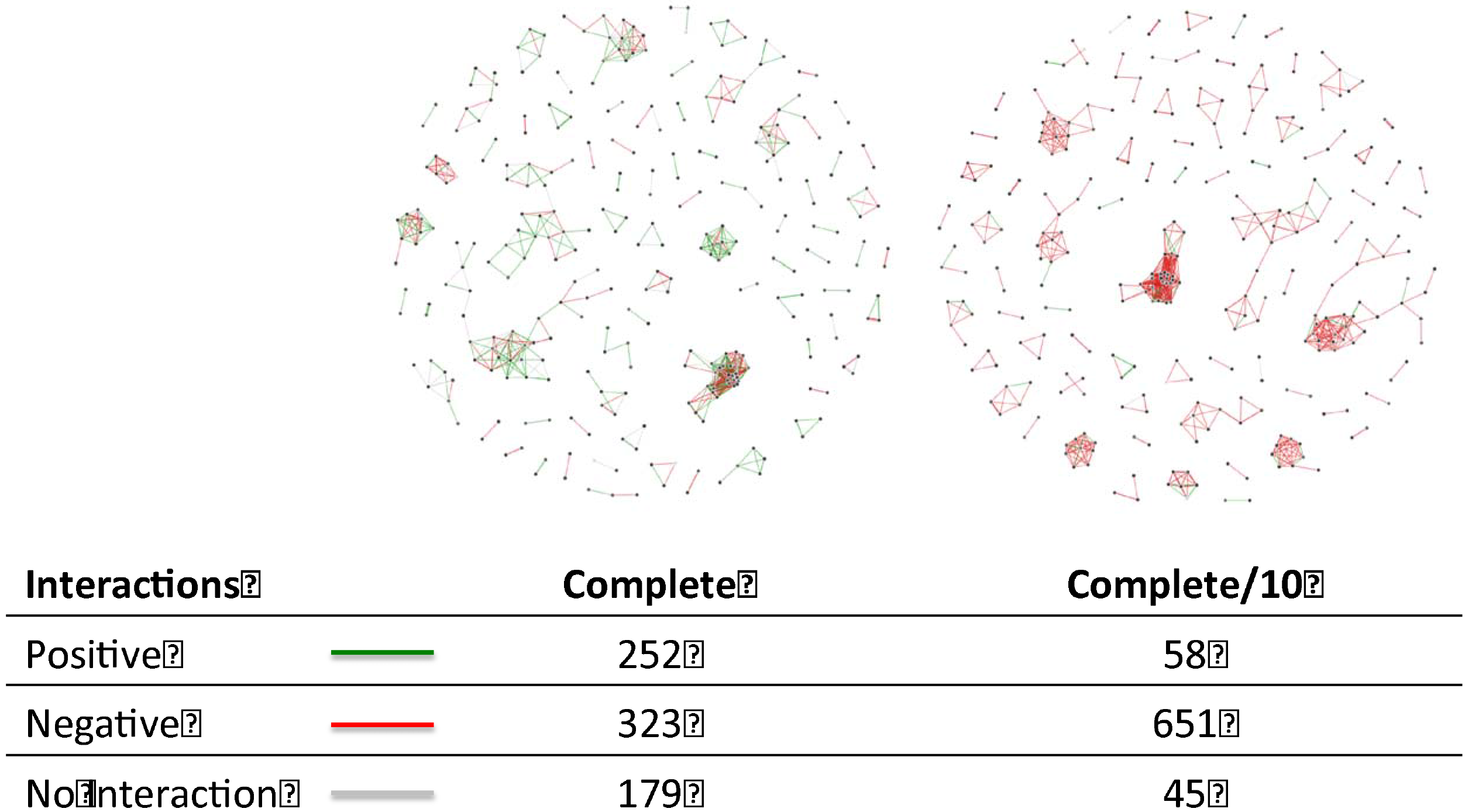
Network and number of the different types of interaction for operational taxonomic units in Case Study **1** under “Complete” (*left*) and “Complete/10” (*right*) metabolic conditions. **There are 380 metabolites in the “Complete” metabolic conditions and they exist as highly available. The metabolic condition “Complete/10” contains the same metabolites as “Complete” but at 10 times lower availability. Please see file Diet.txt for a complete list of the metabolites, and their availabilities represented as uptake metabolic fluxes. It can be seen from the figure that a 10 fold reduction in metabolite availability resulted in a significant decrease in the number of positive interactions predicted to occur between the members of this community, with parallel increase in the number of negative interactions**.

## Results and Discussion

### Usage

Below we present two case studies that exemplify how MMinte can be used to predict microbial interactions under user defined metabolic conditions. All files used in these examples, and a full tutorial on how to perform these analyses, can be found in the Supplementary Materials and in the project folder at github.com/mendessoares/MMinte.

#### Case Study 1

With just two user-provided files, one containing correlations between pairs of OTUs and the other representative sequences from a microbiome study, MMinte creates an association network in which the color of the links represents the types of predicted interactions between pairs of OTUs in a microbial community.

In Case Study 1, we use data from the Human Microbiome Project to demonstrate MMinte's potential for exploratory research studies that focus on the dynamics of host-associated microbial communities. The Human Microbiome Project is a multi-institutional collaboration and the study was reviewed by each participating institution's Institutional Review Boards. Full information can be found in [5]. The data used in this example are a subset of what is available on the Human Microbiome Project page (http://hmpdacc.org/HMQCP/, uncompressed files from rep_set_v13.fna.gz and 0tu_table_v13.txt.gz) [4, 5], allowing users to rerun the analysis in a straightforward and fast way, while taking advantage of publicly available data. The reduced dataset contains 659 associations for 308 OTUs representing 176 species.

**The problem**. The number of positive and negative interactions in a community influences its level of stability and consequently its resistance to invasion by pathogens [43]. With MMinte, we can run our analysis in a variety of metabolic conditions and quantify the number of positive and negative interactions predicted for each. This will generate hypothesis regarding the metabolic conditions likely to favor stability of the system.

**The results**. We ran the full MMinte pipeline by clicking the “run all” tab on MMinte's introductory page and providing two files, corrs.txt (which contains the associations between OTUs) and seqs.txt (which contains representative sequences). Using the default setting of “complete” for the metabolite availability condition, which represents a condition with 380 metabolites available in large amounts. MMinte predicted 252 positive and 323 negative interactions between pairs of OTUs in addition to 179 pairs lacking any type of of interaction (**Figure 2, *left panel***). We then reran Widget 5 with different metabolite availability conditions. **Figure 2** (***right panel***) shows the same correlation network with a metabolite availability that is 10 times lower, resulting in different predicted interactions (58 positive and 651 negative interactions between OTUs and 45 pairs lacking any interaction). This result is consistent with the prediction that lower nutrient availability will favor more competition between organisms.

The results of this analysis highlight some possible characteristics of the community that could not be inferred solely from association data. For instance, assuming stability, and thus protection against pathogen invasion, is greater in communities with more competitive interactions [43], then the metabolic conditions that lead to the predicted network shown in **Figure 2** (***right panel***) are likely to promote more stability. In addition, if we assume that stronger positive correlation values between pairs of species indicate positive interactions [44], the network of interactions observed under metabolic conditions equivalent to the ones listed under “complete” are more reflective of the real system than the alternative metabolic conditions tested. These are just two examples of the window microbe-microbe interactions—the building blocks of community networks—provide for understanding their ecology.

#### Case Study 2

In the following example, we use data from Rey et al [45], who investigated the growth of the sulfate-reducing bacterium *Desulfovibrio piger* in the guts of gnotobiotic mice, in the presence of eight other bacterial species and under different nutritional conditions. *D. piger* is the most commonly found sulfate-reducing bacterium in healthy adults, and is thought to shape the responses of the gut microbiota to dietary changes [45]. However, relatively little is still known about the niche this species occupies and how it may influence the metabolism of the other microbial species found in the gut [45].

**The problem**. The interactions between *D. piger* and other members of the gut microbial community have been shown to influence the level of H_2_S in the gut. However, *D. piger* has a variety of potential metabolic pathways, only some of which will lead to the production of H_2_S. The role of interactions in determining the metabolic niche of *D. piger* in the gut is both important and not fully understood. Using, MMinte, we explored the types of interactions that are predicted to occur between nine different species of microbes that co-occur in the human gut and whose interactions are believed to be metabolically based [45, 46]. We created a set of metabolic conditions where we varied the availability of oxygen, chondroitin sulfate, and fructose. These represent some of the metabolites that were manipulated in the experiments of Rey et al. [45].

**Results**. We started by providing a list of species IDs to Widget 3 (*D. piger*. model 411464.8, *Bacteroides thetaiotaomicron*: model 226186.12, *Bacteroides caccae*: model 411901.7, *Bacteroides ovatus*: model 28116.7, *Eubacterium rectale*: model 657318.4, *Marvinbryantia formatexigens*: model 478749.5, *Collinsella aerofaciens*: model 411903.6, *Escherichia coli*: model 83333.113, and *Clostridium symbiosium*: model 742740.3). After reconstructing the individual species metabolic models and creating two-species communities (*Widget 4*), we predicted species growth rates in the presence and absence of another species in the community by running Widget 5 under 17 different metabolic conditions, listed in Supp. Table 1. To parallel our analysis in Case Study 1, we also calculated the number of positive and negative interactions under metabolic conditions containing 380 metabolites with different availabilities.

A look at the predicted growth rate of *D. piger* in the presence and absence of other species in the community shows that this species is likely to benefit from the presence of each of the other species in the community under “Complete” metabolic conditions. *D. piger* is consistently predicted to grow under aerobic conditions, but under anaerobic conditions, growth is only predicted to occur if either 6. *ovatus, B. thetatiotaomicron, B. caccae, C. symbiosium*, or *E. coli* are present. Thus, using the models reconstructed using ModelSEED, MMinte predicts an obligate association between *D. piger* and these species in anaerobic environments (**Supp. Table 1**). Interestingly, *D. piger* impaired the growth of most species it was paired with under all conditions except “complete”(**Supp. Table 1**). Exceptions were *E. coli* and *E. rectale-*, the magnitude of the effect of *D. piger* on their growth depended on the flux conditions for oxygen, chondroitin sulfate, and sulfate (**Supp. Table 1**). Even though our analysis only focused on variations in three metabolites, the results provide some insight into the niches that these species occupy and they are predicted to interact under a variety of metabolic conditions

Overall, the number of each type of interaction changed depending on metabolite availability, but not linearly (**Figure 3**). For instance, with a lo-fold decrease in metabolite availability, the number of predicted parasitic interactions increased—but with a further lo-fold decrease in metabolite availability, the number of predicted parasitic interactions then decreased. This suggests that alternative metabolic pathways may be invoked depending on the amount of particular metabolites and not necessarily on their presence or absence, affecting how different organisms interact with each other. These results are in concordance with the observation that the nutrient conditions that organisms experience are predicted to have marked effects on the kinds of interactions they have (**Figure 3**).

**Figure 3.**
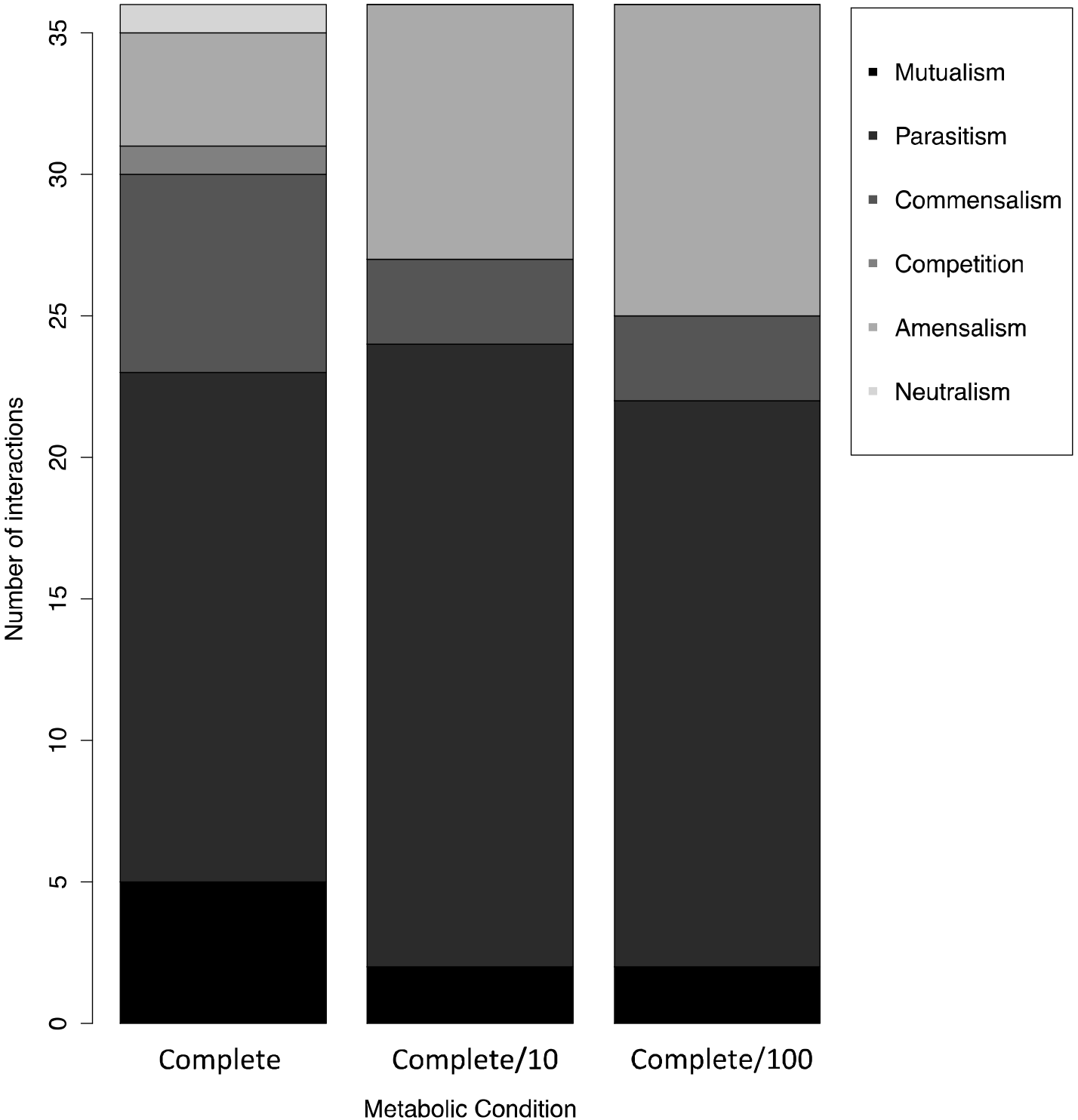
Number of each type of interaction predicted to occur between pairs of the nine bacterial species (*D. piger, Bacteroides thetaiotaomicron, Bacteroides caccae, Bacteroides ovatus, Eubacterium rectale, Marvinbryantia formatexigens, Collinsella aerofaciens, Escherichia coli, and Clostridium symbiosium*) inoculated into the guts of gnotobiotic mice under different metabolic conditions in [45] and used in Case Study 2. The metabolic conditions simulated in MMinte were “Complete”, “Complete/10” and “Complete/100”. There are 380 metabolites in the “Complete” metabolic conditions and they exist as highly available. The metabolic condition “Complete/10” contains the same metabolites as “Complete” but at 10 times lower availability and “Complete/100” contains the same metabolites as “Complete” but at 100 times lower availability. Please see file Diet.txt for a complete list of the metabolites, and their availabilities represented as uptake metabolic fluxes.

**General Discussion**. MMinte bridges the gap between association and mechanism in microbe-microbe interactions by assessing the metabolic influence that two microbes have on each other. Our predictive modeling approach involves reconstructing the pairwise metabolic community models that make up the basic unit of interaction within a community. More specifically, MMinte advances microbiome research by assigning functional interactions instead of simply calculating associations or correlations based on abundance [44]. This allows us to capture the effect of metabolite exchange on the interactions of an entire microbial community across different nutrient conditions, thus providing an important link to the overall drivers of environmental dynamics.

The metabolic interactions that MMinte identifies can be used to understand the broader ecological features of a biological system. Dynamic ecological features such as stability and robustness are linked to competitive-cooperative interactions and the nature of the positive-negative feedback loops they engender [43]. For example, it has been widely posited that negative interactions “self-regulate” and stabilize fluctuations within a community [47, 48]. In Case Study 1, MMinte showed that out of 754 total associations detected among a subset of human microbiome species, 33.4% were predicted to be positive and 42.8% negative under “complete” metabolic conditions. The rest (23.7%) were predicted to not represent significant metabolic interactions between the species. When fasting conditions are modeled by decreasing the availability of metabolites by an order of magnitude, MMinte predicts that 7.7% of interactions will be positive, 86.3% will be negative, and in 6% of the cases, no interactions will occur. This finding intuitively matches the expectation that competition increases in a community with limited nutrient availability [49]. MMinte enables users to grasp these important ecological interactions and better understand the role of competition and cooperation in community stability [50, 51].

Case Study 2 highlights the modular nature of MMinte and the user's ability to explore the effect that changes in the availability of a particular metabolite may have on the interactions between organisms. The results give us an important window into the role of the environment, specifically the presence or absence of oxygen, chondroitin sulfate, and sulfate, on the interactions between *D. piger* and other organisms commonly found in the gut. *In vivo* experiments have shown that mice colonized solely with *D. piger* have significantly increased levels of H_2_S, which is a genotoxic metabolite that may be involved in the development of colorectal cancer [52, 53], compared to mice colonized with a consortium containing the other eight species analyzed here. Understanding the metabolite conditions favoring the dominance of the other species over *D. piger* can help inform dietary interventions aimed at reducing the abundance of this species in the gastrointestinal tract.

Like all algorithms, MMinte has potential limitations. For example, MMinte's predictions are only as accurate as the metabolic models used. These metabolic models are linked from 16S in a multi-step process that involves identification of genomes and metabolic network reconstruction using ModelSEED [28]. Missing data in the genome database or in the biochemical database are both potential sources of error. Conversely, as databases rapidly grow, so will the accuracy of MMinte's predictions. MMinte helps the user minimize the potential for over-interpretation by visually displaying the percent similarity between 16S rDNA provided by the user and the genomic data,

## Conclusions

MMinte is the first tool that predicts the type of interactions occurring between organisms in a complex microbial community under defined metabolite conditions based on the metabolic models of each species. A full run takes data of an association network and 16S rDNA sequences, identifies the genomes, reconstructs metabolic models, and estimates the effect of being in a two-species community for each species under user defined metabolic conditions. The predicted interactions are then plotted in an interaction network. Additionally, the widgets that make up MMinte can be run independently allowing the user to run specific tasks and bypass some of the steps of the analysis. We have incorporated the design principles of clear modularity, usability, and open access into the development of MMinte. In our view, part of the value of MMinte to the development of predictive community metabolic modeling is the potential for integration into other analytical platforms. We view the ability to build on existing development efforts as critical to expanding systems biology tools to wider and broader scales of ecology and data [14]. MMinte is thus a fundamental tool for exploring a large number of interactions, allowing researchers to move beyond the use of statistical measures of association into biologically relevant analysis of interactions between the species in a microbiome.

## Availability of supporting data

All files required to reproduce the results are provided in the project folder at http.github.com/mendessoares/MMinte.

Additionally, results and supplementary files can be found in the following project folder **MMinte/supportFiles/ResultsAndSupplMaterial**. The folder **ResultsCaseStudy1** contains output files from the first example workflow. The folder **ResultsCaseStudy2** contains the file with the Supplementary Table 1, listing the growth rates of the 9 species analyzed in the second example in the presence and absence of another species in the two-species communities under a variety of metabolic conditions.

## Ethics statement

In Case Study 1, we use data from the Human Microbiome Project to demonstrate MMinte's potential for exploratory research studies that focus on the dynamics of host-associated microbial communities. The Human Microbiome Project is a multi-institutional collaboration and the study was reviewed by each participating institution's Institutional Review Boards. Full information can be found in [5].

## Abbreviations

MMinte - Microbial Metabolic interactions

FBA - Flux Balance Analysis

COBRA - Constraint-Based Reconstruction and Analysis

## Author contributions

HMS, NC and MM designed the algorithm. HMS, MM and LMS implemented the algorithm. HMS performed the case studies' analyses. HMS and NC wrote the manuscript. All authors read and approved the final manuscript.

## Acknowledgements

The authors thank all members of the Chia Laboratory, particularly Dr. Vanessa Hale, for testing the software, discussions and critical reading of the manuscript. This work was supported by the Mayo Clinic Center for Individualized Medicine and the National Institutes of Health under award number R01CA179243.

